# Age- and Sex-Specific Fear Conditioning Deficits in Mice Lacking Pcdh10, an Autism Associated Gene

**DOI:** 10.1101/2020.05.25.115634

**Authors:** Sarah L. Ferri, Holly C. Dow, Hannah Schoch, Ji Youn Lee, Edward S. Brodkin, Ted Abel

## Abstract

*PCDH10* is a gene associated with Autism Spectrum Disorder. It is involved in the growth of thalamocortical projections and dendritic spine elimination. Previously, we characterized mice *Pcdh10* haploinsufficient mice *(Pcdh10^+/−^* mice) and found male-specific social deficits that are rescued by N-methyl-D-aspartate receptor (NMDAR) partial agonist d-cycloserine, increased ultrasonic vocalizations in pups, and dark phase hypoactivity. In addition, we determined that the basolateral amygdala (BLA) of these mice exhibited increased dendritic spine density of immature morphology, decreased NMDAR expression, and decreased gamma synchronization. Here, we further characterize *Pcdh10^+/−^* mice by testing for fear memory, which relies upon BLA function. We used both male and female *Pcdh10^+/−^* mice and their wild-type littermates at two ages, juvenile and adult, and in two learning paradigms, cued and contextual fear conditioning. We found that males at both ages and in both assays exhibited fear conditioning deficits, but females were only impaired as adults in the cued condition. These data are further evidence for male-specific alterations in BLA-related behaviors in *Pcdh10^+/−^* mice, and suggest that these mice may be a useful model for dissecting male specific brain and behavioral phenotypes relevant to social and emotional behaviors.

## 1. Introduction

Protocadherin 10 (Pcdh10) is a calcium-dependent transmembrane cell adhesion molecule belonging to the δ2 subfamily of nonclustered protocadherins in the cadherin superfamily (Hirano et al., 1999; Nakao et al., 2008; Kim et al., 2011). PCDH10 (OL-protocadherin) is located on chromosome 4 in humans and chromosome 3 in mice and is expressed almost exclusively in the mammalian brain, particularly in the striatum, amygdala, and cerebellum (Hirano et al., 1999; Aoki et al., 2003; Kim et al., 2011). Pcdh10 has been associated with cell migration (Nakao et al., 2008), thalamocortical and corticothalamic projections (Uemura et al., 2007), and proteasomal degradation of PDS-95 to promote synapse elimination (Tsai et al., 2012).

Genome-wide analyses and loci mapping have revealed copy number variations in the *PCDH10* gene and its regulatory region in association with Autism Spectrum Disorder (ASD), a neurodevelopmental disorder that causes deficits in communication and social interaction and repetitive or restricted behaviors (Morrow et al., 2008; Bucan et al., 2009; Taylor et al., 2020). Because ASD has been linked to a large number of genes encoding synaptic cell adhesion molecules, many transgenic mouse models have been created, including a *Pcdh10* knockout model in which the first exon was replaced with a lacZ-neo selection cassette (Uemura et al., 2007; Taylor et al., 2020).

Because mice homozygous for *Pcdh10* deletion do not survive past 3-4 weeks of age, likely due to failure of striatal axon outgrowth, we recently characterized heterozygous mice (Pcdh10^+/−^) (Uemura et al., 2007). We found that male and female *Pcdh10^+/−^* pups at postnatal day 6 emitted increased maternal separation-induced ultrasonic vocalizations compared to their WT littermates. We also uncovered a male-specific deficit in juvenile social approach behavior that is rescued with systemic administration of d-cycloserine, a partial NMDAR agonist at the glycine site. In addition, we demonstrated that these mice have no evidence of anxiety-like behavior in elevated zero maze and intact olfactory habituation-dishabituation despite strong Pcdh10 expression in the olfactory bulb. Female mutants had decreased motor performance in rotarod compared to their wild-type littermates. *Pcdh10^+/−^* mice did not exhibit any deficits in novel object recognition task and no repetitive behaviors were observed (Schoch et al., 2017). Males, but not females demonstrated hypoactivity in long-term home-cage activity monitoring specifically in the dark phase (Angelakos et al., 2019). Interestingly, *Pcdh10^+/−^* males had a number of changes in structure and function of the basolateral amygdala (BLA). Using voltage sensitive dye, we showed that *Pcdh10^+/−^* males exhibited reduced power of gamma band activity in LA-BLA transmission compared to WT littermates, indicating reduced connectivity between these areas. In addition, *Pcdh10^+/−^* males exhibited increased spine density of immature morphology (filopodial spines) on LA/BLA pyramidal neurons and decreased expression of GluN1A and GluN2A NMDAR subunits in the postsynaptic fraction of BLA cells compared to those of WT littermates (Schoch et al., 2017).

*Pcdh10^+/−^* males are known to have abnormalities in the BLA and deficits in social behaviors that engage the BLA (Ferri et al., 2015; Schoch et al., 2017). This raises the question of whether *Pcdh10^+/−^* males might have deficits in fear conditioning, a behavior that involves the BLA (Ressler and Maren, 2019), and that is disrupted in several in mouse models relevant to ASD (Han et al., 2012; Banerjee et al., 2016; Kim et al., 2016), and in autistic humans (Gaigg and Bowler, 2007; Top Jr. et al., 2016). We hypothesized that fear conditioning would be disrupted in *Pcdh10^+/−^* mice, and we tested juvenile and adult male and female Pcdh10^+/−^ and WT littermate mice in both contextual and cued fear conditioning paradigms.

## 2. Materials and Methods

### 2.1 Animals

*Pcdh10^+/−^* males were produced by Lexicon Pharmaceuticals, Inc. (Uemura et al., 2007). The first exon of *Pcdh10* was replaced with a *lacZ-neo* selection cassette. Founders were backcrossed with C57BL/6J females for more than 15 generations. Mice were bred and experiments took place both at University of Iowa and University of Pennsylvania. In both cases, *Pcdh10^+/−^* males were crossed with C57BL/6J females to produce heterozygous Pcdh10^+/−^ and wild-type offspring, which were housed with same-sex littermates 2-5 per cage in a temperature- and humidity-controlled environment on a 12-hour light/dark cycle. All mice had access to food and water ad libitum. Behavior was conducted during the light cycle. All mice were cared for in accordance with the National Institutes of Health Guide for the Care and Use of Laboratory Animals and all procedures were approved by the Institutional Animal Care and Use Committees at University of Iowa and University of Pennsylvania.

### 2.2 Cued Fear Conditioning

Five to seven days prior to training, mice were habituated to single housing. Mice were handled for 2-3 min per day for 3-5 days prior to conditioning. On the day of training, each mouse was placed in a chamber with electrified grid flor inside a sound attenuating box (Med Associates, Inc.) for 3 min. After 2 min of free exploration, a tone (2900 Hz, 70 dB) sounded, which lasted 30 s and co-terminated with a 2 s, 1.5 mA shock, and the mouse was removed 30 s after that. Twenty-four hours later, a test session was conducted. The mice were placed in the same training chamber which had been significantly altered into a novel context for 6 min. White plastic inserts covered the electrified grid floor and the walls of the chamber had been altered to change the texture and shape of the compartment from a cube to a cylinder. Several drops of lemon dish soap were placed within the chamber to alter the smell. Freezing behavior was measured continuously by FreezeScan software (CleverSys Inc.).

### 2.3 Contextual Fear Conditioning

Mice were handled and housed as described above. On training day, individual mice were placed in the training chamber described above for 3 minutes. The first 2:28 of free exploration were considered “Baseline,” after which a single 1.5 mA footshock was delivered, lasting 2 s. The mice then remained in the chamber for an additional 30 s. Twenty-four hours later (test session) the mice were placed in the same, unaltered chamber for 5 min. Freezing behavior was measured continuously by FreezeScan software (CleverSys Inc.).

### 2.4 Statistics

For all graphs, repeated measures two-way ANOVAs were used to determine main effects of session (training versus testing), genotype *(Pcdh10^+/−^* vs wild-type), and an interaction. Bonferroni was used as a post hoc test. Significance was set to p<0.05.

## 3. Results

### 3.1 Cued Fear Conditioning

In juvenile male mice undergoing cued fear conditioning (28-32 d), a RM two-way ANOVA revealed a main effect of session (training, Pr-CS and during CS at 24 hr test; F_(2,50)_=38.620, p<0.0001), no main effect of genotype *(Pcdh10^+/−^* vs wild-type; F_(1,25)_=0.529, p=0.474), and a significant genotype x session interaction (F_(2,50)_=6.176, p=0.004). A Bonferroni post hoc test indicated that *Pcdh10^+/−^* males spent a similar percentage of time freezing compared to their wild-type littermates during the baseline period of training (p>0.999), and during the Pre-CS period of the 24 h test, prior to the presentation of the tone in the altered context (p=0.326). However, during the presentation of the tone (CS) during the 24 h test, *Pcdh10^+/−^* males spent significantly less time freezing than their WT littermates (p=0.048; **Fig. 1A**). For juvenile females (28-32 d), a RM two-way ANOVA revealed no main effects of session (F_(2,52)_=59.940, p<0.0001) or genotype (F_(1,26)_=0.005, p=0.942), and no genotype x session interaction (F_(2,52)_=0.004, p=0.996). Bonferroni post hoc analysis showed that *Pcdh10^+/−^* females spent a similar percentage of time freezing compared to their wild-type littermates during all sessions (Baseline, Pre-CS, and CS; p>0.999 for each; **Fig. 1B**).

**Figure 1.**
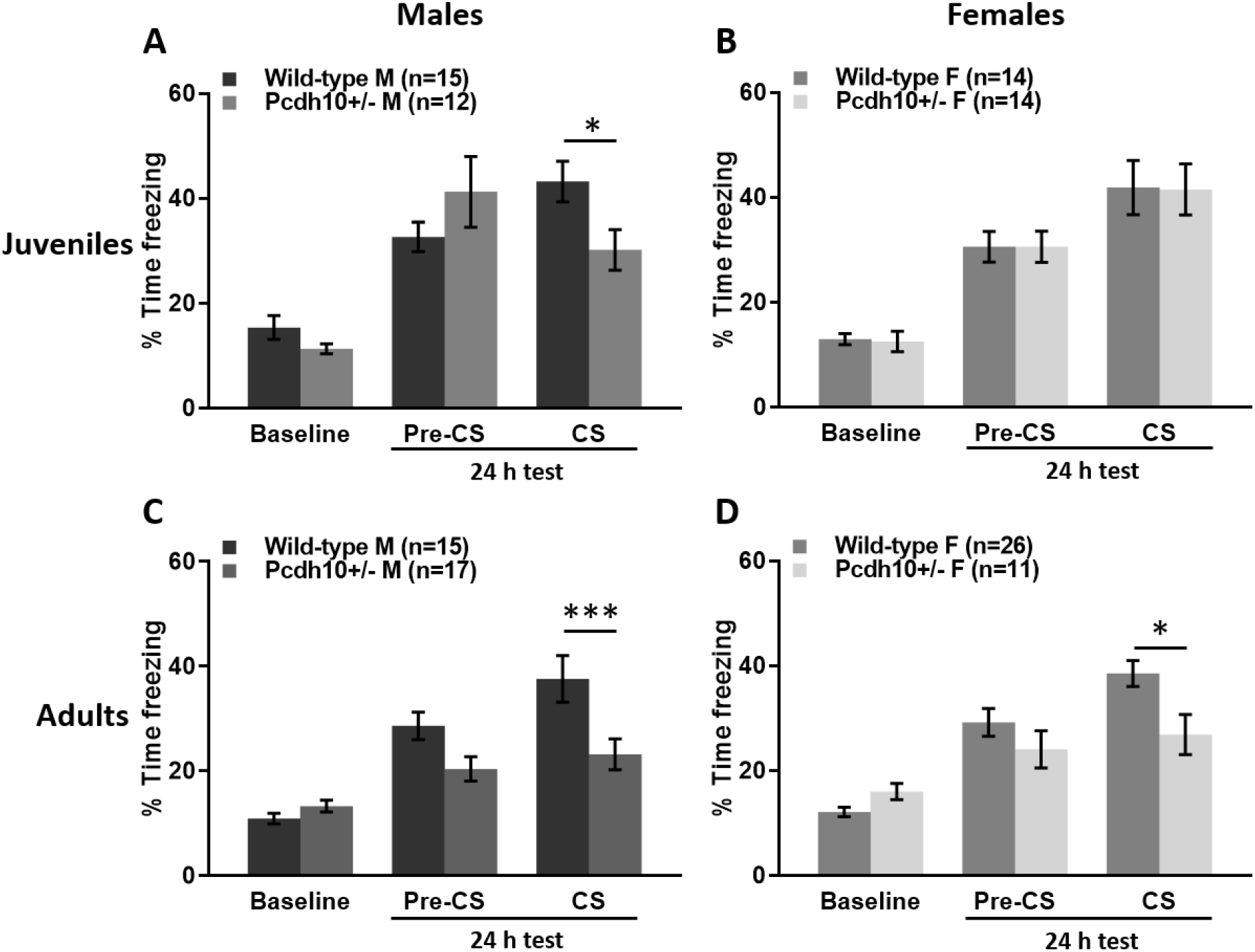
Juvenile and adult males and adult females lacking one copy of *Pcdh10* exhibit cued fear conditioning deficits. A) Juvenile *Pcdh10^+/−^* males spent a similar percentage of time freezing as their wild-type littermates during a baseline period of training and in the pre-CS period, prior to the tone cue during the 24 hr test. During the presentation of the tone cue, however, *Pcdh10^+/−^* males froze less than WT. B) Juvenile *Pcdh10^+/−^* females froze for a similar percentage of time as their wild-type littermates during the baseline period of training and the Pre-CS and CS periods of the 24 h cued fear conditioning test. C) Adult *Pcdh10^+/−^* males spent significantly less time freezing to a tone cue 24 h after training than their WT littermates. Freezing levels were not significantly different during baseline. D) *Pcdh10^+/−^* adult females had similar freezing behavior during a baseline period of training and 24 h later in the Pre-CS period of the cued fear conditioning test. During the CS presentation of the 24 h test, *Pcdh10^+/−^* females froze less than their wild-type littermates. ***=p<0.001, *=p<0.05.

For adult male mice (80-100 d), a RM two-way ANOVA revealed a main effect of session (F_(2,60)_=40.410, p<0.0001), a main effect of genotype (F_(1,30)_=5.516, p=0.026), and a significant genotype x session interaction (F_(2,60)_=8.315, p=0.0006). A Bonferroni post hoc analysis demonstrated that *Pcdh10^+/−^* males spent a similar percentage of time freezing compared to their wild-type littermates during the baseline period of training (p>0.999). *Pcdh10^+/−^* males seemed to freeze less than WT during the Pre-CS period of the 24 h test, however the difference was not significant (p=0.093). *Pcdh10^+/−^* males spent significantly less time freezing during the CS tone during the 24 h test compared to WT littermates (p=0.0007; **Fig. 1C**). Freezing behavior of adult females (80-100 d) in cued fear conditioning was analyzed using a RM two-way ANOVA. Results indicated a main effect of session (F_(2,70)_=39.670, p<0.0001), no main effect of genotype (F_(1,35)_=2.011, p=0.165), and a significant genotype x session interaction (F_(2,70)_=6.683, p=0.002). A Bonferroni post hoc test showed that *Pcdh10^+/−^* females spent a similar percentage of time freezing compared to their wild-type littermates during the baseline period of training (p=0.952) and during the Pre-CS period of the 24 h test (p=0.570). However, during the presentation of the tone (CS) during the 24 h test, they spent significantly less time freezing than their WT littermates (p=0.011; **Fig. 1D**).

In summary, both juvenile and adult *Pcdh10^+/−^* males exhibit deficits in cued fear conditioning compared to their wildtype littermates, while adult, but not juvenile, female *Pcdh10^+/−^* mice show an impairment in cued fear conditioning.

### 3.2 Contextual Fear Conditioning

In juvenile male mice (28-32 d) trained with a single shock and tested in the same context, a RM two-way ANOVA revealed a main effect of session (training vs test; F_(1,24)_=82.010, p<0.0001), a trend towards a main effect of genotype *(Pcdh10^+/−^* vs wild-type; F_(1,24)_=3.570, p=0.071), and a significant genotype x session interaction (F_(1,24)_=5.416, p=0.029). A Bonferroni post hoc test showed that *Pcdh10^+/−^* males spent a similar percentage of time freezing compared to their wild-type littermates during the baseline period of training (p>0.999), but significantly less during the 24 h test (p=0.009; **Fig. 2A**). For juvenile female mice (28-32 d) undergoing contextual fear conditioning, a RM two-way ANOVA revealed a main effect of session (F_(1,20)_=58.570, p<0.0001), and no main effect of genotype (F_(1,20)_=1.375, p=0.255) and no significant genotype x session interaction (F_(1,20)_=3.393 p=0.080). A Bonferroni post hoc test indicated that *Pcdh10^+/−^* females spent a similar percentage of time freezing compared to their wild-type littermates during the baseline period of training (p>0.999). *Pcdh10^+/−^* females seemed to spend less time freezing during the 24 h test, but this difference did not reach statistical significance (p=0.081; **Fig. 2B**).

**Figure 2.**
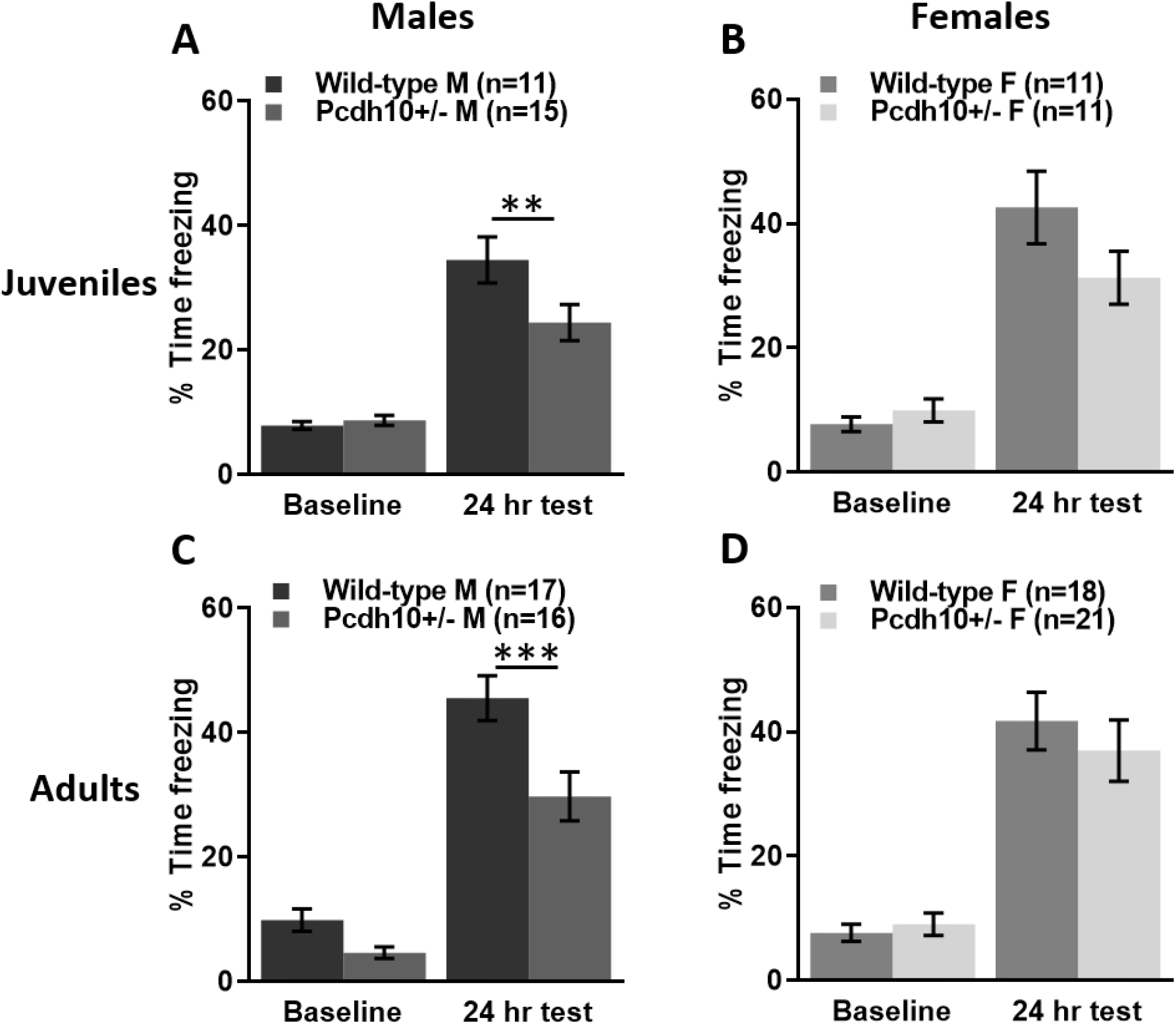
Juvenile and adult males lacking one copy of *Pcdh10* exhibit contextual fear conditioning deficits. A) Juvenile *Pcdh10^+/−^* males spent significantly less time freezing 24 hours after a single-shock contextual fear conditioning training than their wild-type littermates. Baseline freezing prior to the shock was similar between groups. B) Juvenile female *Pcdh10*^+/−^mice seemed to freeze less than wildtype littermates in a long-term memory test of contextual fear conditioning, but this difference was not statistically significant. Baseline freezing was similar between the groups. C) Adult *Pcdh10^+/−^* and WT littermate males spent a similar percentage of time freezing during the baseline period of contextual fear conditioning but *Pcdh10^+/−^* males spent significantly less time freezing 24 h later. D) Adult *Pcdh10^+/−^* females had similar freezing behavior compared to their wild-type littermates, both at baseline and at the 24 h test. ***=p<0.001, **=p<0.01.

For contextually-conditioned adult male mice (80-180 d) receiving a single shock during training and tested in the same context 24 h later, a RM two-way ANOVA revealed a main effect of session (training vs test; F_(1,31)_=207.300, p<0.0001), a main effect of genotype *(Pcdh10^+/−^* vs wild-type; F_(1,31)_=9.329, p=0.005), and a significant genotype x session interaction (F_(1,31)_=6.249, p=0.018). A Bonferroni post hoc test demonstrated that *Pcdh10^+/−^* males spent a similar percentage of time freezing compared to their wild-type littermates during the baseline period of training (p=0.401), but significantly less during the 24 h test (p=0.0005; **Fig. 2C**). For adult female mice (80-180 d) undergoing contextual fear conditioning, a RM two-way ANOVA revealed a main effect of session (F_(1,37)_=99.150, p<0.0001), no main effect of genotype (F_(1,37)_=0.174, p=0.679), and no genotype x session interaction (F_(1,37)_=0.967, p=0.332). A Bonferroni post hoc test showed that *Pcdh10^+/−^* females spent a similar percentage of time freezing compared to their wild-type littermates during the baseline period of training (p>0.999) and during the 24 h test (p=0.709; **Fig. 2D**).

In summary, both juvenile and adult *Pcdh10^+/−^* males demonstrate contextual fear conditioning deficits compared to their wildtype littermates, while *Pcdh10^+/−^* females at both ages are unaffected.

## 4. Discussion

Fear conditioning has been strongly associated with BLA function. BLA lesions disrupt both cued and contextual fear conditioning (LeDoux et al., 1990; Goosens and Maren, 2001). Contextual fear conditioning also involves the hippocampus, particularly the dorsal portion and CA3 (Clark and Squire, 1998). We have previously characterized several abnormalities in the LA/BLA of Pcdh10^+/−^ males, including an increase in density of filopodial dendritic spines (Schoch et al., 2017). These spines are immature and likely have decreased synaptic functioning (Harris, 1999) and may prevent the plasticity necessary for fear conditioning (Rogan et al., 1997; Nabavi et al., 2014). In addition, we showed decreased gamma synchronization in the LA-BLA circuit (Schoch et al., 2017). Gamma oscillations are known to be important for fear memory retrieval and can modulate BLA neuronal firing (Popescu et al., 2009; Stujenske et al., 2014; Bocchio et al., 2017). Finally, we demonstrated a decrease in NMDAR expression in the BLA of *Pcdh10^+/−^* males (Schoch et al., 2017). NMDAR function is important for fear memory; it has been shown that NMDAR antagonist infusion into the BLA blocked fear conditioning acquisition (Campeau et al., 1992). While hippocampus of *Pcdh10^+/−^* mice has not been thoroughly investigated, behavior in another hippocampus-dependent task, the Morris water maze, is intact in these mice (unpublished). Further investigation of the role of the hippocampus will be important.

There are other important players in fear memory learning as well, including BLA GABAergic interneurons (Bocchio et al., 2017). Neurotransmitters such as dopamine, noradrenaline, and acetylcholine are also important (de la Mora et al., 2010; Heath et al., 2015; Kwon et al., 2015; de Oliveira et al., 2017; Wilson and Fadel, 2017; Giustino and Maren, 2018; Brandão and Coimbra, 2019; Stubbendorff et al., 2019; Tang et al., 2020). Serotonin and glutamate influence fear memory (Walker and Davis, n.d.; Maren, 1996; Bauer, 2015; Johnson et al., 2015; Bocchio et al., 2016), as does the transcription factor CREB (cyclic AMP response element-binding protein) (Vazdarjanova and McGaugh, 1998; Bocchio et al., 2017; Ressler and Maren, 2019). These important elements have not yet been investigated in *Pcdh10^+/−^* mice. Similarly, structure and function of other brain areas in *Pcdh10^+/−^* mice are not well studied. For example, the central amygdala is necessary for fear learning as well (LeDoux et al., 1990; Wilensky et al., 2006).

Interestingly, as with social deficits, *Pcdh10^+/−^* males preferentially exhibited fear conditioning deficits (Schoch et al., 2017). Both juvenile and adult males displayed deficits in both cued and contextual fear conditioning, whereas females exhibited deficits only as adults in the cued paradigm. This may lend support to the theory that females require higher genetic burden to present ASD-associated phenotypes than males (Jacquemont et al., 2014; Ferri et al., 2018). Although we have determined that *Pcdh10^+/−^* females have unaltered spine density in the BLA (Schoch, unpublished), we have not investigated NMDAR expression or LA-BLA connectivity. It may be that these or other factors are disrupted in females, or slightly abnormal such that behavioral deficits are only manifest under specific conditions such as in cued fear conditioning, or only in older females, whereas males display deficits at all ages in both memory paradigms. This robust sex difference is interesting and requires further investigation. Exploring these mechanisms may shed light on the male bias of neurodevelopmental disorders.

Like *Pcdh10^+/−^* mice, several mouse models associated with ASD exhibit fear conditioning deficits, including CD38, Homer1a, and Scn1a mutants (Han et al., 2012; Banerjee et al., 2016; Kim et al., 2016), as well as a number of others (Markram et al., 2008; Stapley et al., 2013; Howell et al., 2017; Nolan et al., 2017; Fricano-Kugler et al., 2019). Reports of fear conditioning in individuals with ASD have been varied, with some reporting impairments, and some failing to find differences compared to controls (Bernier et al., 2005; Gaigg and Bowler, 2007; Sterling et al., 2013). Recent findings have indicated that amygdala response to fear conditioning is attenuated in individuals with ASD compared with age-matched controls (Top Jr. et al., 2016). In addition, changes in amygdala volume, cell size, and responsivity have been reported in individuals with ASD (Bauman and Kemper, 1985; Abell et al., 1999; Baron-Cohen et al., 2000; Critchley et al., 2000; Sparks et al., 2002; Barnea-Goraly et al., 2004; Schumann and Amaral, 2006; Conturo et al., 2008; Pinkham et al., 2008; Kohls et al., 2012; Kleinhans et al., 2016), indicating that this brain area is an important player in the neurobiology of ASD. Our present results demonstrating cued and contextual fear conditioning deficits in *Pcdh10^+/−^* males indicate that fear may be a readout of the function of circuits that are impacted in autism.

## Acknowledgements

This work was supported by the National Institutes of Health Grants 1P50MH096891 (Raquel Gur)-subproject 6773 (ESB and TA), R01MH080718 (ESB), K01MH119540 (SF), and Pennsylvania Department of Health (SAP #4100042728) (to Robert Schultz, TA, ESB). TA was supported by the Brush Family Chair of Biology at the University of Pennsylvania. TA and SLF are supported by the Roy J. Carver Charitable Trust at the University of Iowa. The content is solely the responsibility of the authors and does not necessarily represent the official views of the National Institute of Mental Health or the National Institutes of Health. The Pennsylvania Department of Health specifically disclaims responsibility for any analyses, interpretations, or conclusions. We would also like to thank the Neural Circuits and Behavior Core at the University of Iowa.

## Notes

### Competing Interest Statement

The authors have declared no competing interest.

